## Supplementary figures and images for "Resolving multisensory and attentional influences across cortical depth in sensory cortices"

### SupplementatyFigure-1

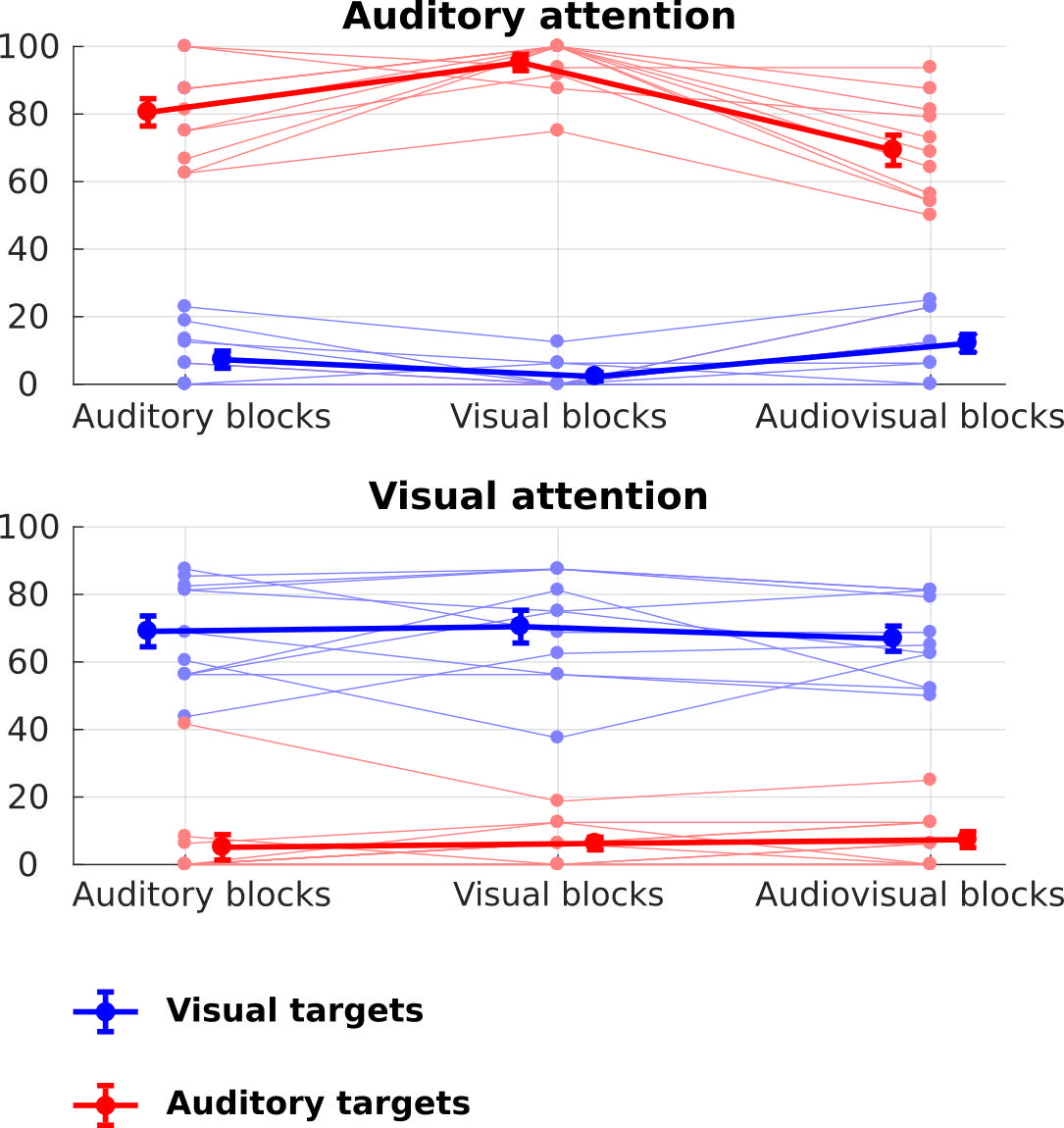

### SupplementatyFigure-2

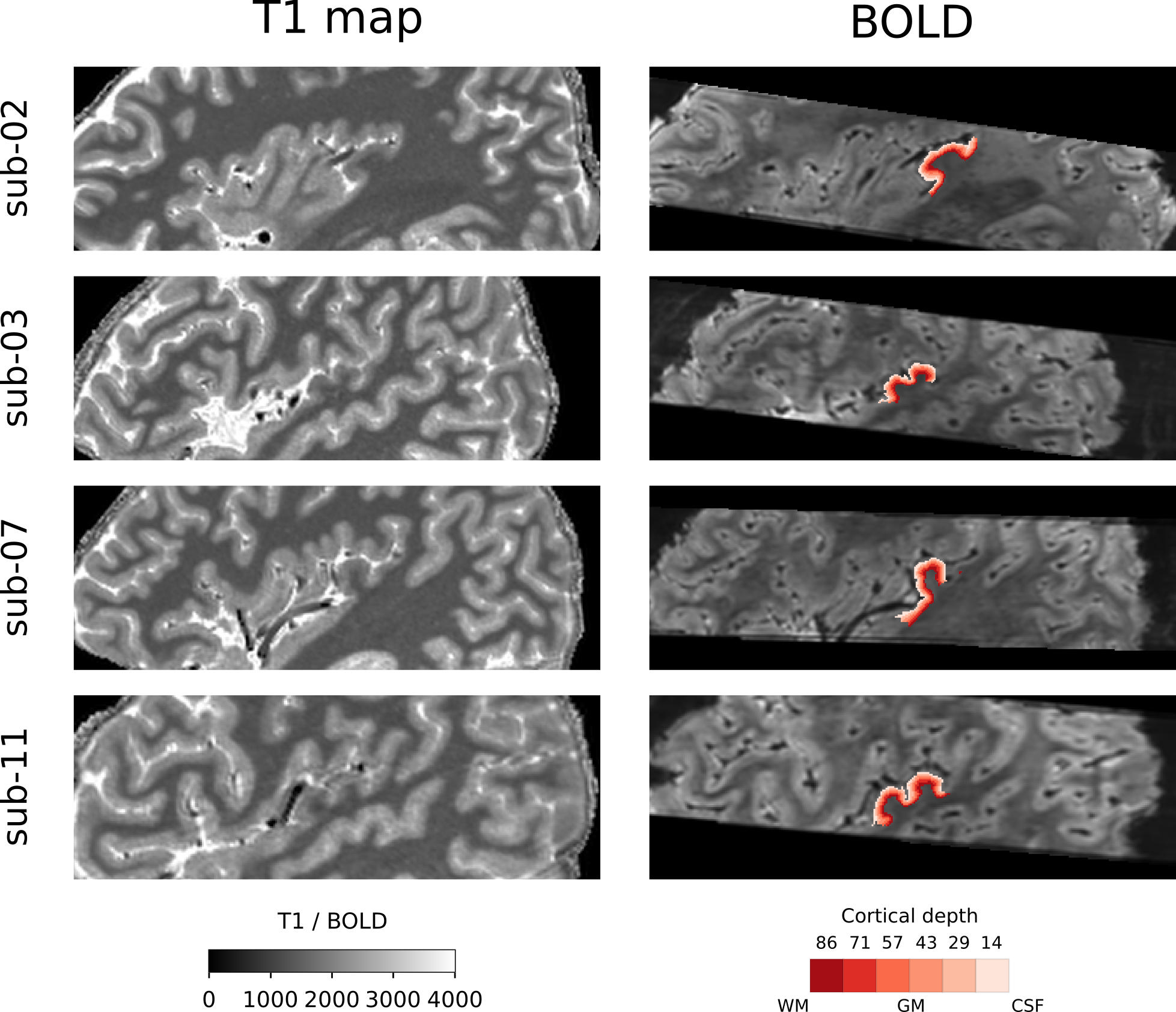

### SupplementatyFigure-3

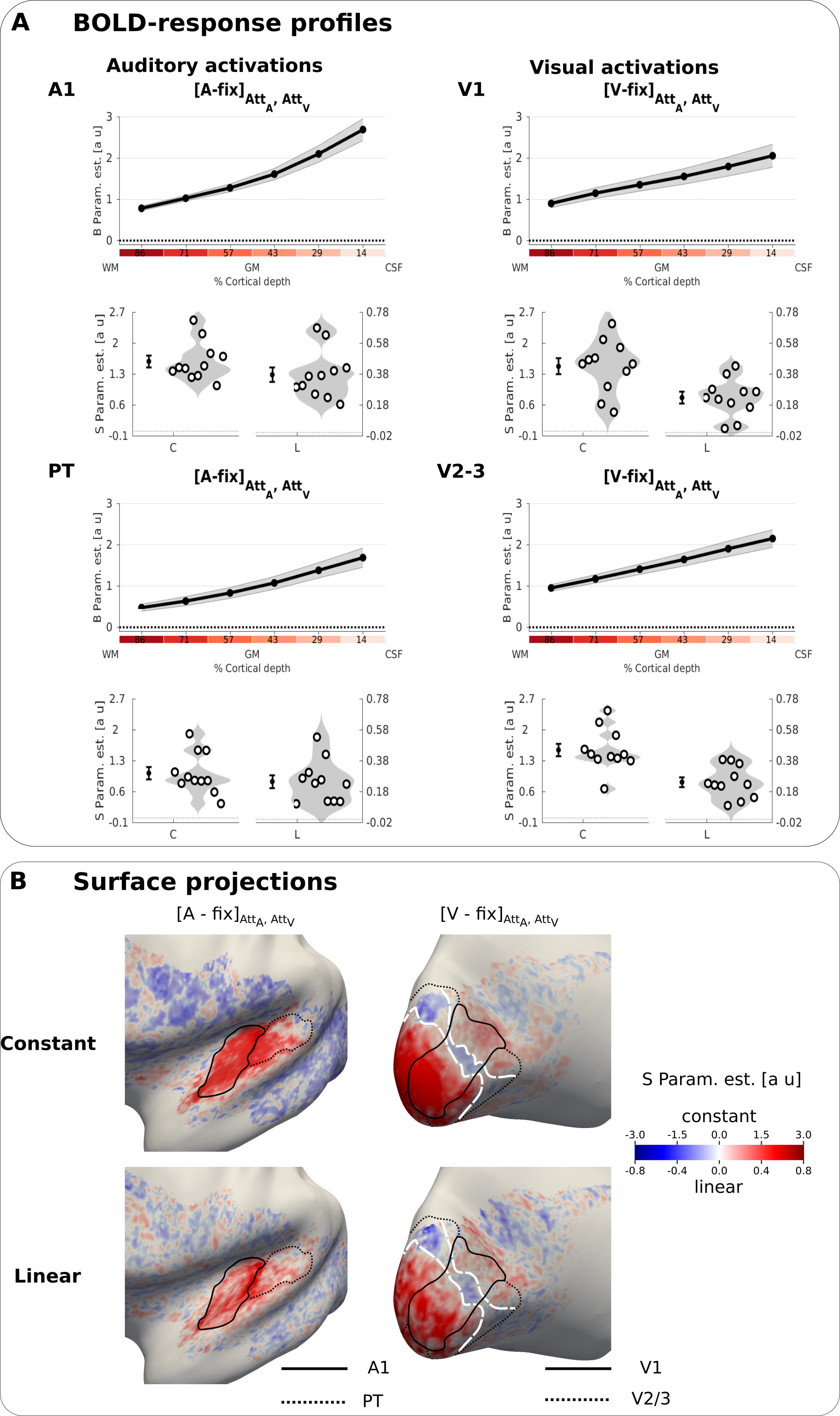

### SupplementatyFigure-4

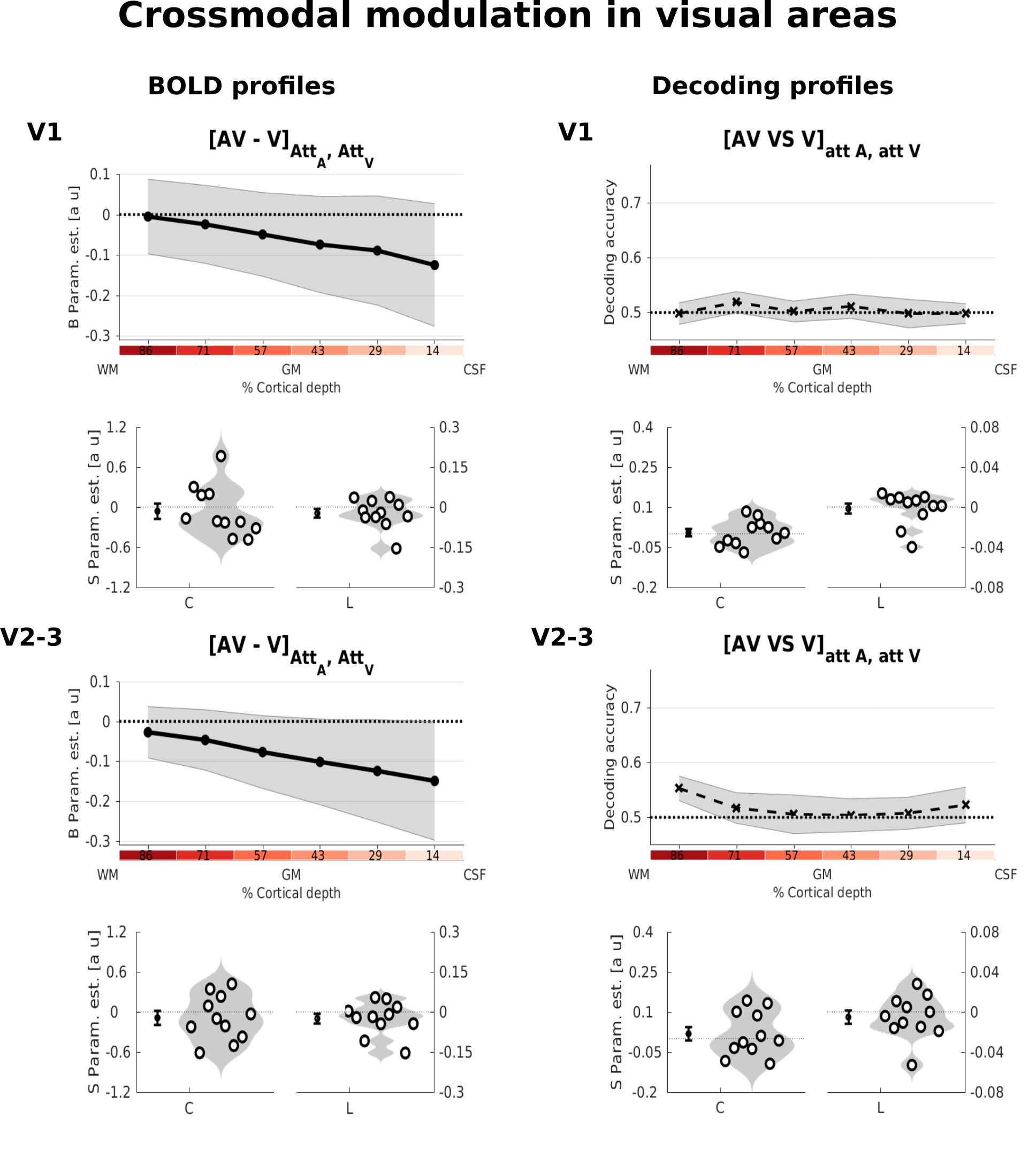

### SupplementatyFigure-5

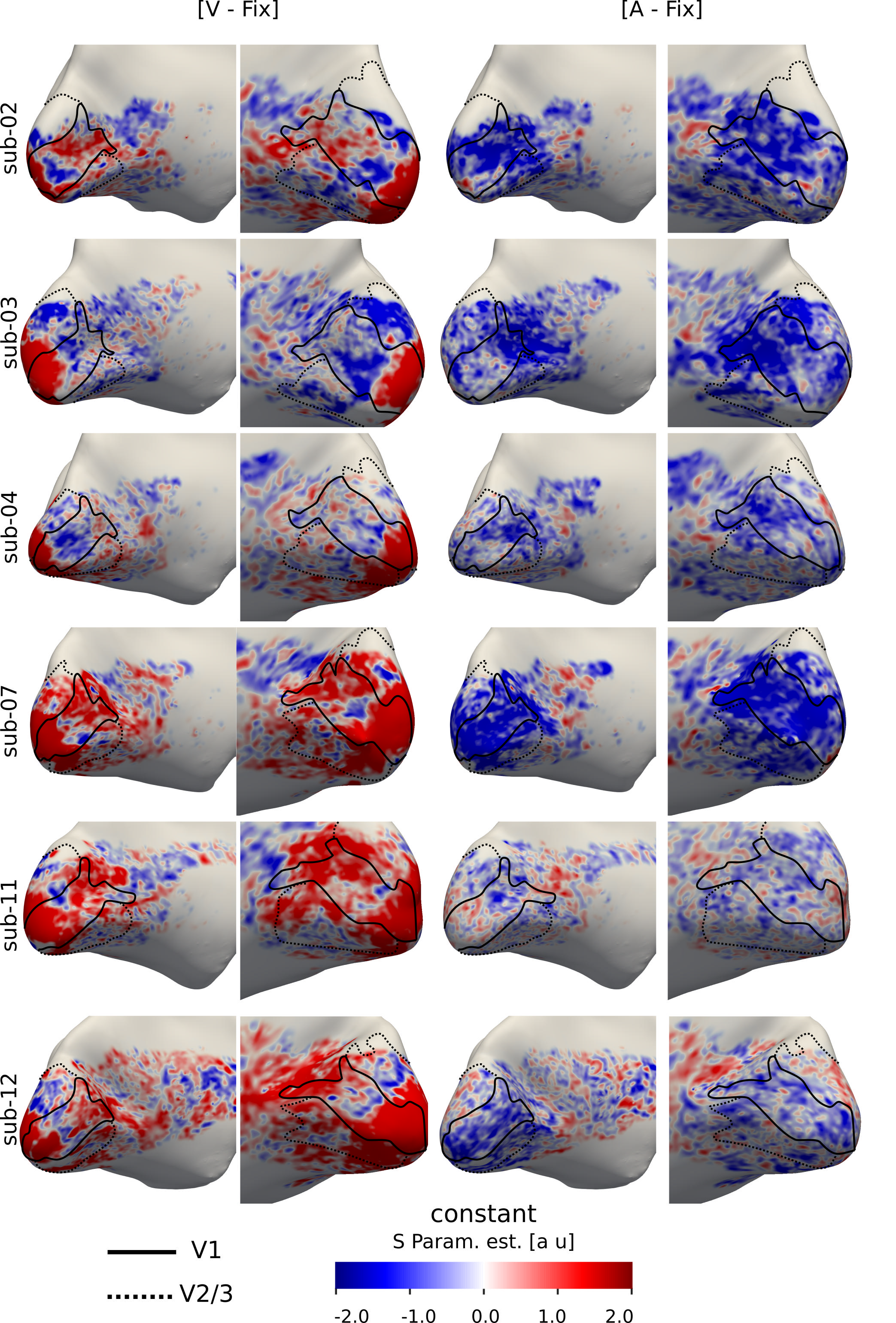

### SupplementatyFigure-6

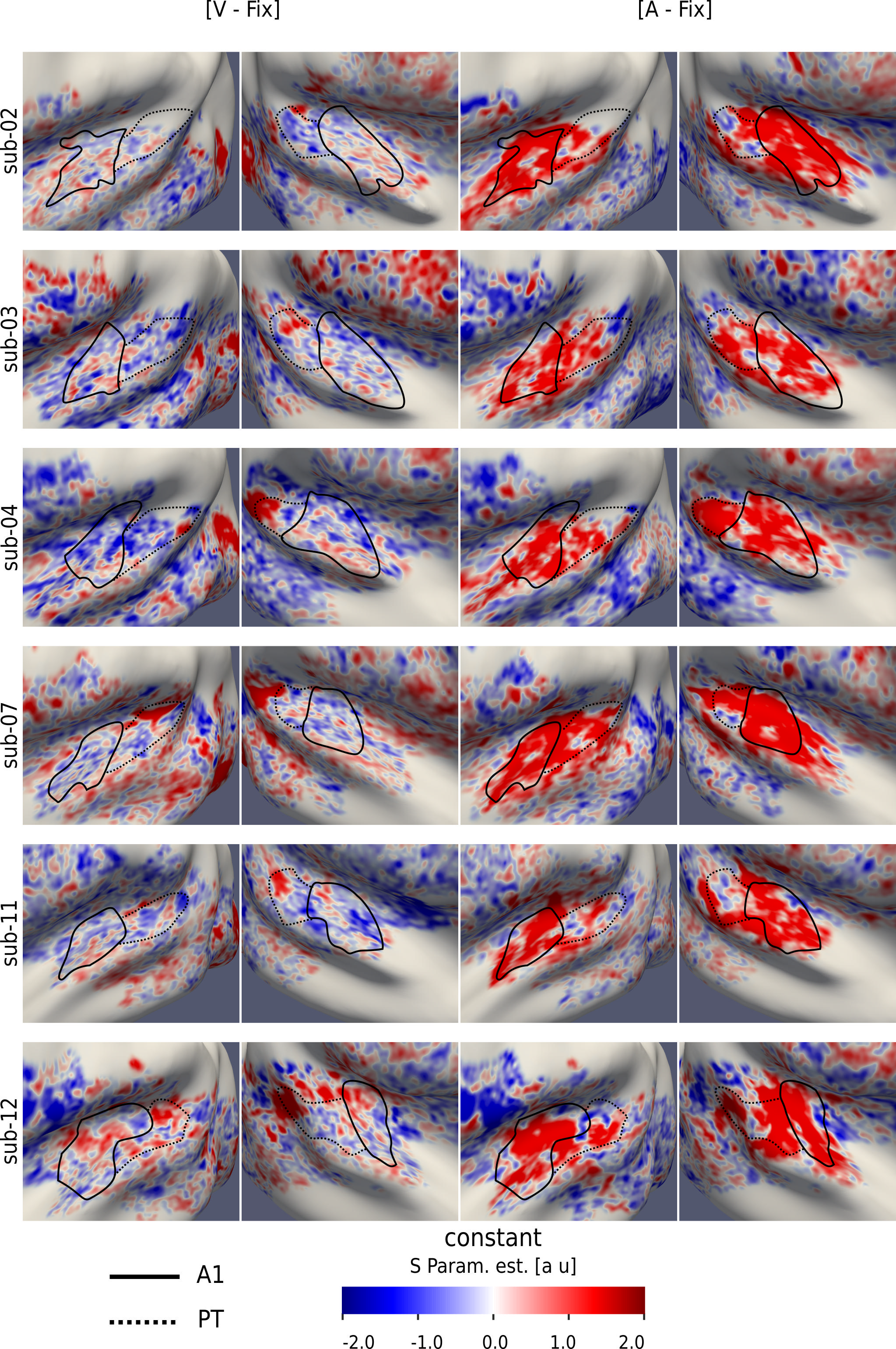

### SupplementatyFigure-7

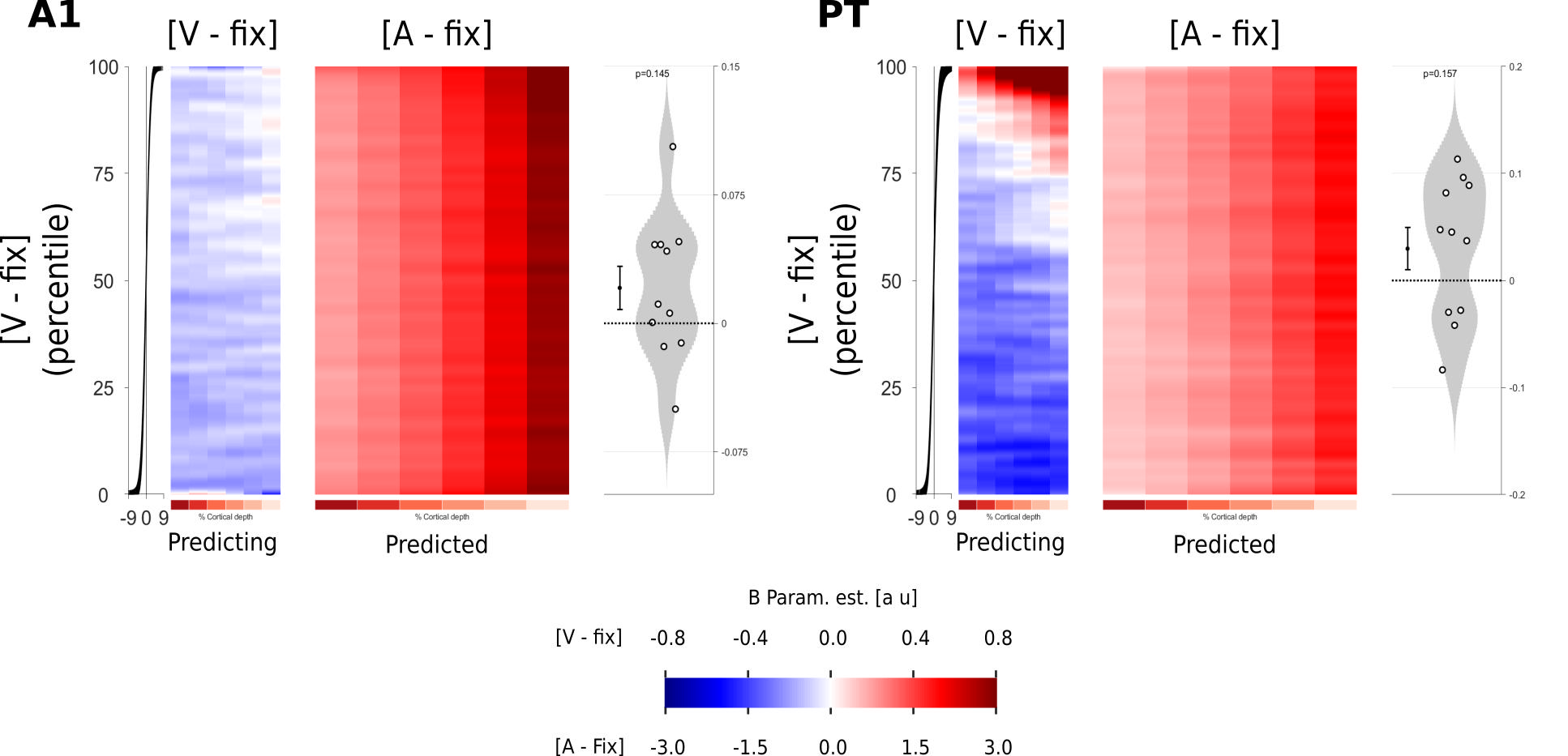
